# *pgxRpi*: an R/Bioconductor package for user-friendly access to the Beacon v2 API

**DOI:** 10.1101/2025.03.20.644282

**Authors:** Hangjia Zhao, Michael Baudis

## Abstract

**Motivation:** The Beacon v2 specification, established by the Global Alliance for Genomics and Health (GA4GH), consists of a standardized framework and data models for genomic and phenotypic data discovery. By enabling secure, federated data sharing, it fosters interoperability across genomic resources. Progenetix, a reference implementation of Beacon v2, exemplifies its potential for large-scale genomic data integration, offering open access to genomic mutation data across diverse cancer types.

**Results:** We present *pgxRpi*, an open-source R/Bioconductor package that provides a streamlined interface to the Progenetix Beacon v2 REST API, facilitating efficient and flexible genomic data retrieval. Beyond data access, *pgxRpi* offers integrated visualization and analysis functions, enabling users to explore, interpret, and process queried data effectively. Leveraging the flexibility of the Beacon v2 standard, *pgxRpi* extends beyond Progenetix, supporting interoperable data access across multiple Beacon-enabled resources, thereby enhancing data-driven discovery in genomics.

**Availability and Implementation:** *pgxRpi* is freely available under the Artistic-2.0 license from Bioconductor (https://doi.org/doi:10.18129/B9.bioc.pgxRpi), with actively maintained source code on GitHub (https://github.com/progenetix/pgxRpi). Comprehensive usage instructions and example workflows are provided in the package vignettes, available at https://github.com/progenetix/pgxRpi/tree/devel/vignettes.

## 1 Introduction

The Beacon v2 specification, adopted as a Global Alliance for Genomics and Health (GA4GH) standard in 2022, provides a standardized framework and data models for secure, federated discovery of genomic and phenotypic data. This addresses critical challenges in data sharing for biomedical research and clinical applications [1], [2]. Compared to Beacon v1, which was primarily designed for existence queries on genomic variant collections, returning only binary “Yes” or “No” responses [3], Beacon v2 significantly expands its scope. It now supports richer queries, enabling detailed retrieval of both genomic variant data and phenotype information, making it more suitable for clinical and translational research. As a reference implementation of Beacon v2 specification, Progenetix—an open-access genomic data resource established in 2001—offers comprehensive mutation profiles of cancer genomes, with a primary focus on copy number variations (CNVs) across diverse human neoplasms [4]. Currently, it hosts approximately 150,000 samples spanning nearly 900 distinct cancer types. These data can be accessed via a user-friendly web graphical user interface (GUI) or programmatically through a representational state transfer (REST) application programming interface (API), fully compliant with the Beacon v2 protocol.

This paper introduces *pgxRpi*, an R/Bioconductor package that simplifies querying and retrieval of phenotypic and variant data from Progenetix through the Beacon v2 REST API. In addition to facilitating data access, *pgxRpi* provides powerful visualization and processing tools to support data exploration and downstream analysis. By leveraging the Beacon v2 standard, *pgxRpi* not only serves as a companion to the Progenetix API but also demonstrates its flexibility for connecting with other “beaconized” resources, underscoring the potential of Beacon v2 in fostering interoperable and efficient genomic data discovery.

## 2 Methods

### 2.1 Retrieving data from the Progenetix Beacon v2 API

The Progenetix Beacon v2 API, accessible at https://progenetix.org/ beacon, facilitates programmatic access to structured cancer genomic data. To streamline data integration in R, we developed the `pgxLoader` function (Figure 1), which converts API responses from JSON format into relational tables, enabling more efficient manipulation and analysis. This conversion process is guided by a YAML configuration file located in the `inst/config` directory of the package, ensuring compliance with the Beacon v2 specification.

**Figure 1.**
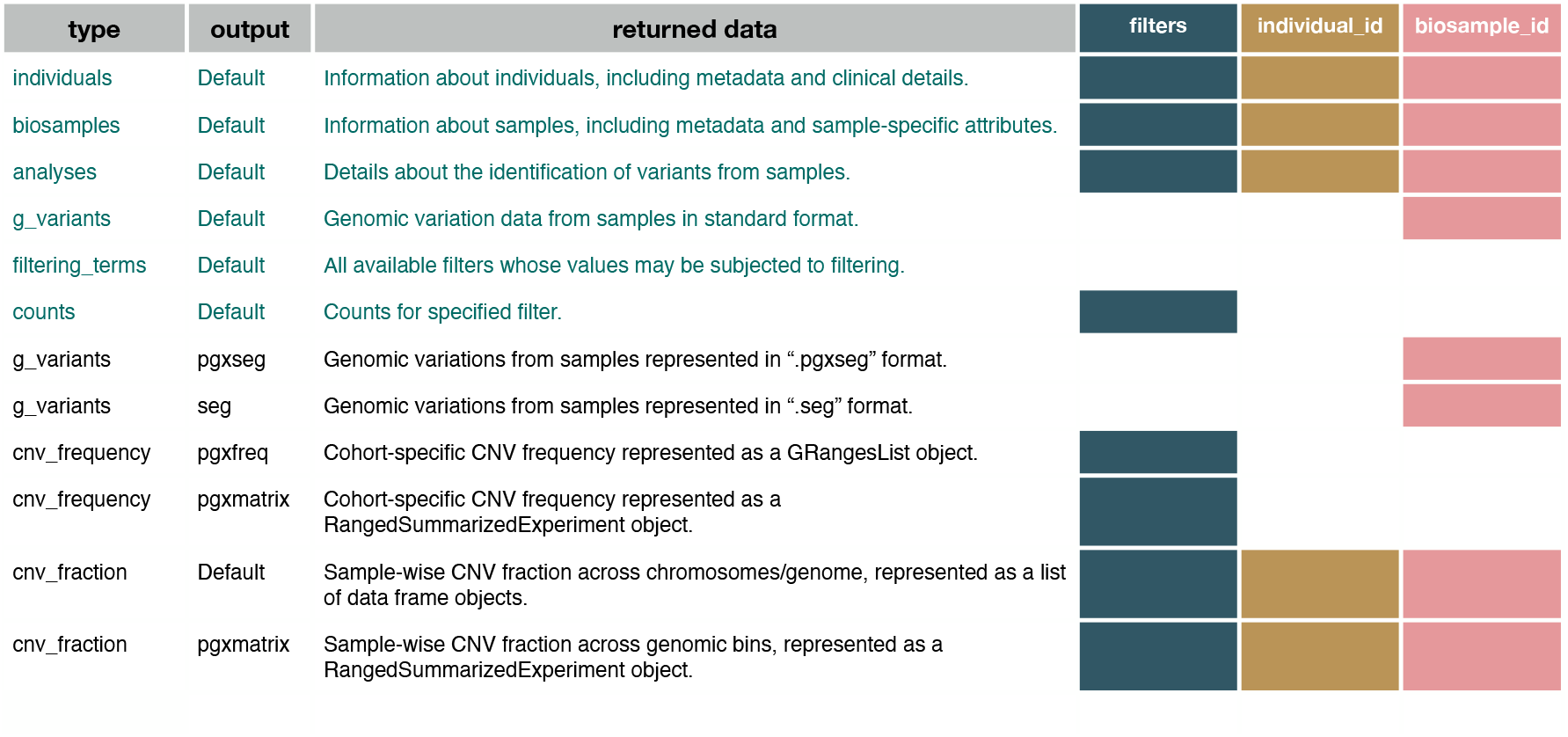
Data retrieval options using the pgxLoader function. The first two columns display the available options for the `type` and `output` parameters of `pgxLoader`, while the third column provides descriptions of the retrieved data. Colored boxes indicate the accessible data types based on the specified search criteria in Progenetix. Data highlighted in green is retrieved through the standard Beacon v2 API, whereas data highlighted in black is accessed via the Progenetix-specific “services” API.

Progenetix data is organized into four main entities in accordance with the Beacon v2 model: individuals, biosamples, analyses, and genomic variations. These correspond to the REST API resources `/individuals`, `/biosamples`, `/analyses`, and `/g_variants`, respectively. Users can query these entities through `pgxLoader` by specifying the `type` parameter. This unified approach eliminates the need to use different endpoints for each data category, simplifying data retrieval^1^.

Refinements to queries can be achieved using additional parameters such as `biosample_id`, `individual_id`, and `filters`. These parameters allow users to filter results based on criteria defined by the Beacon v2 standard. For instance, `biosample_id` and `individual_id` correspond to unique identifiers for biosamples and their associated individuals (*cf*. `/biosamples/`*{*`id`*}* and `/individuals/`*{*`id`*}* in REST format), while `filters` define rules for selecting records based on specific field values, predominantly referencing bio-ontologies or registered identifiers through the use of compact uniform resource identifiers (CURIEs).

To aid in constructing queries, the Beacon v2 specification includes a `/filtering_terms` informational endpoint, which lists all available fields and their filterable values. This functionality is integrated into the `pgxLoader` function by setting the `type` parameter accordingly, enabling efficient and precise querying. By default, the Beacon v2 specification applies AND logic when multiple filters are specified, ensuring more specific query results. The example below demonstrates how to retrieve sample details using NCI Thesaurus (NCIt) [5] ontology terms as filters, specifically selecting biosamples from male (C20197) lung adenocarcinoma (C3512) patients:

~~~
pgxLoader(type=“biosamples”, filters=c(“NCIT:C3512”,”NCIT:C20197”))
~~~

In addition to retrieving detailed records, `pgxLoader` also supports count queries, which return the number of results matching specified filters. This feature is accessible by setting `type=“counts”`, providing a quick overview of data availability before executing full queries.

### 2.2 Data retrieval from Beacon v2 compliant resources

The Beacon v2 specification comprises two key components: the framework and the data models. The framework standardizes request and response formats, enabling consistent communication between clients and data providers, while the data models define the structure for representing biological data. This architecture supports seamless data access across diverse genomic resources aligned with the Beacon v2 model.

The `pgxLoader` function leverages this architecture to query data from any Beacon-compliant resource by specifying the `domain` and `entry_point` parameters. For instance, details of individuals with severe COVID-19 infections (C189227) can be retrieved from the Genomic Data Infrastructure (GDI) Spain Node as follows:

~~~
pgxLoader(type=“individuals”, filters=“NCIT:C189227”,
domain=“https://beacon-spain.ega-archive.org“, entry_point=“api”)
~~~

The Beacon framework ensures standardized communication, although variations in data representation may occur due to differences in data types, access policies, and data granularity across Beacon instances. To address these challenges, *pgxRpi* employs a YAML-based mapping system that harmonizes Beacon v2 responses into a unified format suitable for downstream analysis. This mapping system is optimized for Progenetix but also performs effectively with resources that share structural similarities, such as *cancercelllines*.*org* [6]. This resource adopts the same middleware and API stack, built using the *bycon* Python package, and adheres to the Beacon v2 API standard.

By leveraging the Beacon v2 framework and data models, *pgxRpi* can efficiently extract and harmonize key data from diverse Beacon-enabled databases, even in the presence of structural differences. This capability not only supports integration of heterogeneous datasets but also ensures the package’s adaptability to future Beacon protocol updates.

Beyond querying individual Beacon instances, *pgxRpi* supports asynchronous multi-domain queries, enabling users to retrieve data from multiple Beacon v2 resources in parallel. The `num_cores` parameter allows users to control the number of processing cores, accelerating query execution and reducing overall processing time. The following example demonstrates how to retrieve the number of available records from female (C16576) patients across the RD-Connect Genome-Phenome Analysis Platform (GPAP) [7], Progenetix, and *cancercelllines*.*org* Beacon v2 nodes:

~~~
pgxLoader(type=“counts”, filters=“NCIT:C16576”,
domain=c(“https://playground.rd-connect.eu“,
“https://progenetix.org“, “https://cancercelllines.org“),
entry_point = c(“beacon2/api”, “beacon”, “beacon”))
~~~

By enabling parallel data retrieval, this functionality optimizes query efficiency and minimizes computational overhead, making it a powerful tool for large-scale data exploration.

### 2.3 Employing the extended functionality of the Progenetix API

The Progenetix Beacon v2 API extends the standard Beacon v2 protocol by providing additional functionality through the “services” endpoint, enabling specialized access to the full extent of Progenetix data. Services can be accessed in R using the same `pgxLoader` function (Figure 1), ensuring a consistent and streamlined approach to querying.

By setting the `type` parameter, users can retrieve additional data entities beyond the standard Beacon v2 model, including “cnv frequency” and “cnv fraction”, which provide pre-calculated CNV features useful for genomic analysis. Additionally, users requiring data formats optimized for analytical workflows can specify the `output` parameter to obtain results in the desired format. This flexibility ensures that retrieved data aligns with specific analytical needs and facilitates downstream processing.

### 2.4 Data visualization and analysis

*pgxRpi* provides a suite of functions designed to enhance the visualization and analysis of genomic data, allowing users to effectively explore and interpret retrieved datasets. For survival analysis, the `pgxMetaplot` function, built on the *survminer* package [8], generates Kaplan–Meier survival plots from queried individual data, facilitating comparisons of survival outcomes across different groups. For CNV studies, the `pgxFreqplot` function, built on the *copynumber* package [9], visualizes CNV frequency data, helping users identify cohort-specific patterns and generate hypotheses. Supplementary Figure S1 presents example outputs of these visualization functions.

Beyond visualization, *pgxRpi* offers utility functions to streamline data processing. The `segtoFreq` function computes CNV frequency from segment variant data, supporting both standard “.seg” files and “.pgxseg”—a Progenetix-specific format that integrates CNV variants with metadata. Additionally, `pgxSegprocess` facilitates the processing of local “.pgxseg” files downloaded from the Progenetix website by extracting segment variants and metadata and organizing them into structured data frames for further analysis. This function also incorporates `pgxMetaplot`, `segtoFreq`, and `pgxFreqplot`, creating a comprehensive toolkit for survival analysis and CNV frequency computation and visualization. By automating key analytical steps, *pgxRpi* simplifies the workflow for Beacon-enabled genomic data analysis.

## 3 Conclusion

The *pgxRpi* package provides a simple, user-friendly interface for accessing and analyzing Beacon v2-compatible genomic data, with Progenetix as a reference implementation. By transforming API responses into structured, analysis-ready formats, *pgxRpi* enhances data accessibility and usability and provides integration with R-based analytical workflows. Typical examples here could be the support of variant calling for CNV analyses through integration of disease-matched CNV frequency data or the comparison of genomic variant calls to reference datasets in clinical genomics applications.

Beyond Progenetix, *pgxRpi* extends to other Beacon v2-compliant resources, facilitating interoperable, federated genomic data exploration. This versatility establishes it as a powerful tool for cross-resource data integration but also underscores the value of the Beacon protocol beyond data discover, as a facilitator of federated data analysis. Here, through the support of GA4GH’s Beacon v2 specification, *pgxRpi* advances standardized and scalable genomic data sharing and fosters collaborative research and innovation in genomics.

## 4 Abbreviations

GA4GH: Global Alliance for Genomics and Health
CNV: Copy number variation
GUI: Graphical user interface
REST: Representational state transfer
API: Application programming interface
CURIE: Compact uniform resource identifier
NCIt: National Cancer Institute thesaurus
GDI: Genomic Data Infrastructure
GPAP: Genome-Phenome Analysis Platform

## Footnotes

1 The additional *runs* entity and its `/runs` endpoint are formally supported by the Progenetix API but provide minimal data content.

## References

[1] H. L. Rehm, A. J. Page, L. Smith, et al., “Ga4gh: International policies and standards for data sharing across genomic research and healthcare,” Cell genomics, vol. 1, no. 2, 2021.

[2] J. Rambla, M. Baudis, R. Ariosa, et al., “Beacon v2 and beacon networks: A “lingua franca” for federated data discovery in biomedical genomics, and beyond,” Human mutation, vol. 43, no. 6, pp. 791–799, 2022.

[3] M. Fiume, M. Cupak, S. Keenan, et al., “Federated discovery and sharing of genomic data using beacons,” Nature biotechnology, vol. 37, no. 3, pp. 220–224, 2019.

[4] Q. Huang, P. Carrio-Cordo, B. Gao, R. Paloots, and M. Baudis, “The progenetix oncogenomic resource in 2021,” Database, vol. 2021, baab043, 2021.

[5] National Cancer Institute, Nci thesaurus, 2025. [Online]. Available: https://ncithesaurus.nci.nih.gov/.

[6] R. Paloots and M. Baudis, “Cancercelllines. org—a novel resource for genomic variants in cancer cell lines,” Database, vol. 2024, baae030, 2024.

[7] S. Laurie, D. Piscia, L. Matalonga, et al., “The rd-connect genome-phenome analysis platform: Accelerating diagnosis, research, and gene discovery for rare diseases,” Human mutation, vol. 43, no. 6, pp. 717–733, 2022.

[8] A. Kassambara, M. Kosinski, and P. Biecek, Survminer: Drawing survival curves using ‘ggplot2’, R package version 0.5.0, 2024. [Online]. Available: https://rpkgs.datanovia.com/survminer/index.html.

[9] G. Nilsen, K. Liestøl, P. Van Loo, et al., “Copynumber: Efficient algorithms for single-and multi-track copy number segmentation,” BMC genomics, vol. 13, pp. 1–16, 2012.

