## Supplementary Figure S1 for "*pgxRpi*: an R/Bioconductor package for user-friendly access to the Beacon v2 API"

### Supplementary figures

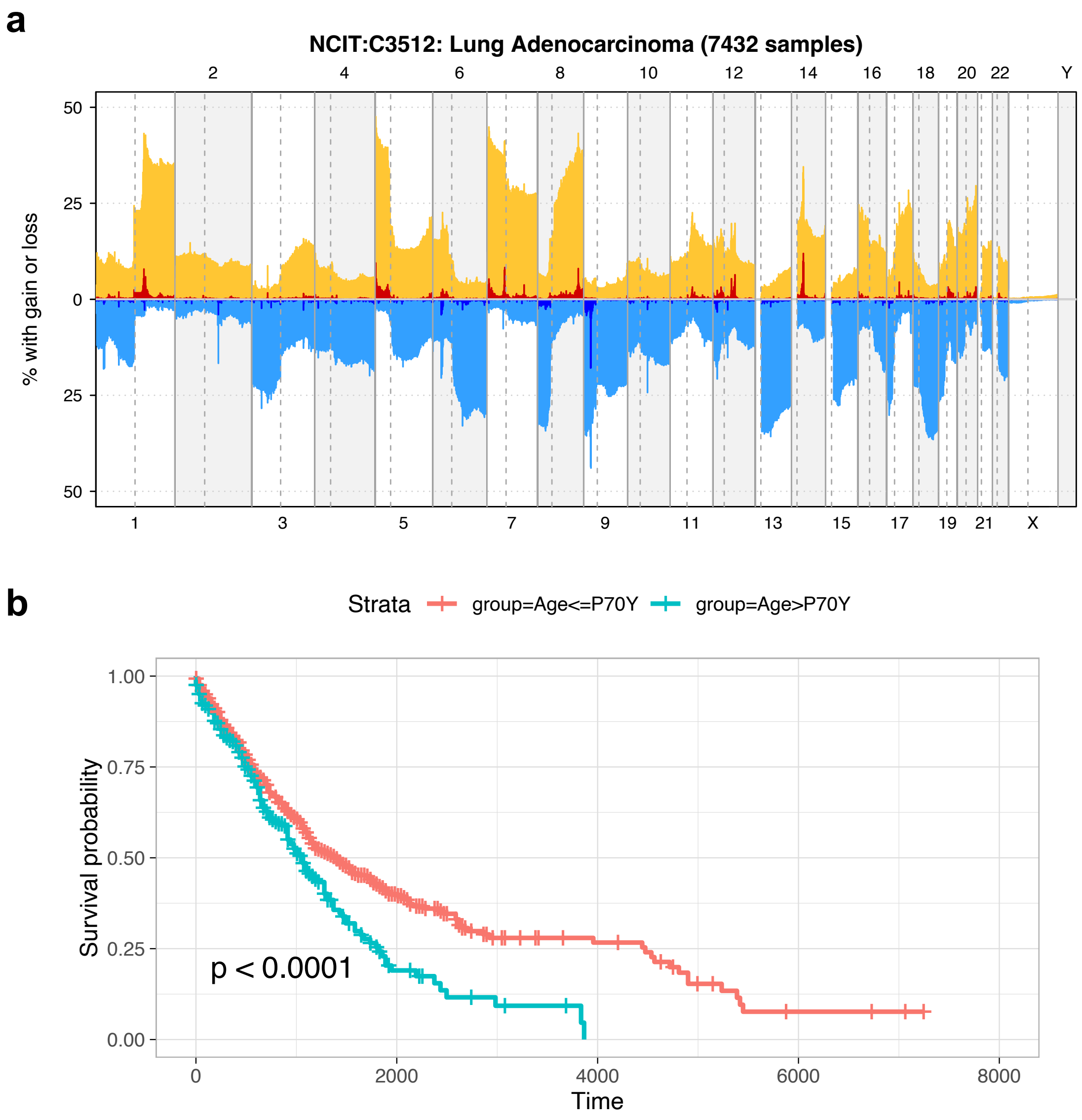


**Supplementary Figure S1: Visualization examples.** (a) Output of the *pgxFreqplot* function. The Y-axis represents the percentage of lung adenocarcinoma samples in Progenetix with CNAs in 1 MB genomic bins, while the X-axis denotes chromosomal positions. Orange and red indicate low-level and high-level duplications, whereas light blue and dark blue represent low-level and high-level deletions, respectively. (b) Output of the *pgxMetaplot* function. A Kaplan–Meier survival plot comparing survival differences between two age groups based on clinical data from lung adenocarcinoma samples in Progenetix.
